# EEG Microstates reveal distinct network dynamics in lucid and non-lucid REM sleep

**DOI:** 10.1101/2025.02.12.637792

**Authors:** Xinlin Wang, Emma Peters, Johannes Strelen, Nathan Lockhart, Michael Franklin, Stephen LaBerge, Thomas Koenig, Martin Dresler, Daniel Erlacher

## Abstract

During lucid dreaming, the dreamer is aware that they are dreaming. In past years, numerous studies have explored the characteristics of lucid dreams, which link lucid dreams to vivid visual perception, positive emotions, and wake-like metacognition and executive functions. To determine brain network activity associated with the conscious experience during lucid dreaming, we analyzed 39 sleep recordings (non-lucid REM vs. lucid REM) with 32-channel polysomnography from 8 lucid dreamers, using EEG microstate analysis. We found that microstates A and G dominated during lucid REM sleep compared to non-lucid REM sleep, and microstates B, C, and D dominated during non-lucid REM sleep compared to lucid REM sleep. We explored the correlation of our microstate maps with previous findings based on topographical similarities of the microstate maps in our study. This suggests that in our study, microstate A might be associated with emotional processing, microstate B with visual processing, microstate C with salience network activity, microstate D with executive functions, and microstate G with the default mode network. Our results suggest that lucidity during REM sleep is associated with increased self-visualization, metacognition, and executive processing, along with decreased emotional processing and reduced default mode network activity. Additionally, we found the inverse relationship between the presence of microstates/networks associated with regions that serve specific functions and evidence for the function being used. This might indicate the inhibitory function of the EEG microstates during sleep. Our study provides novel insight into the distinct network dynamics in lucid and non-Lucid REM sleep.

## Introduction

During sleep, we perceive a complex dream world with all our senses. These dream experiences integrate bizarre images and themes into a simulated world. They are characterized by rich primary consciousness (Hobson, 2009), and deficits in the secondary consciousness (Hobson, 2009). Dreamers usually lack insight into the fact that they are dreaming and experience limited thought (e.g., rare self-awareness) (Domhoff, 2023; Kahan & LaBerge, 2011). Lucid dreaming is an exception in this regard: A lucid dream is a dream during which the dreamer is aware that they are dreaming (Kahan & LaBerge, 1994). During lucid dreaming, dreamers are able to signal their realization of dreaming using specific eye movement patterns, and even completion of a prearranged task (Dresler et al., 2011; Erlacher et al., 2014). In the past years, numerous studies have explored the characteristics of lucid dreams with different methods. Studies applying questionnaires or dream reports suggested that lucid dreams might be linked to increased positive emotions (Schredl, 2024; Voss et al., 2013), perception hallucinations (Drinkwater et al., 2020), metacognition (Dresler et al., 2014; Drinkwater et al., 2020; Stumbrys et al., 2015; Voss et al., 2013) and highly attentional ability (Blagrove et al., 2010; Stumbrys & Erlacher, 2017). By using EEG, researchers indicated that the power in the beta frequency band (13–19 Hz) was increased during lucid REM in both parietal regions (Holzinger et al., 2006) and gamma band (40Hz) in frontal and frontolateral regions (Voss et al., 2009) compared to non-lucid REM sleep. In neuroimaging studies, compared to non-lucid REM sleep, lucid REM sleep showed increased BOLD activity in the right dorsolateral prefrontal cortex and bilateral frontopolar areas (Dresler et al., 2012a). The frontopolar cortex gray matter volume was positively associated with experienced dream lucidity and metacognitive monitoring (Filevich et al., 2015). In addition to the localized effects observed in the specific brain regions, a study by Baird et al. demonstrated that individuals reporting frequent lucid dreaming exhibit increased functional connectivity between the anterior prefrontal cortex (aPFC) and temporoparietal association areas (Baird et al., 2018). This pattern of functional connectivity is associated with increased cognitive control and self-awareness. In light of this, it is particularly interesting to investigate whether lucid REM sleep is associated with the activation of specific global brain networks compared to non-lucid REM sleep.

EEG microstates have emerged as a promising tool for measuring the spontaneous fluctuations of activity in large-scale brain networks underlying different conscious experiences (Michel & Koenig, 2018). Researchers have attempted to link microstates to specific underlying neural networks to interpret these variations. In a recent meta-analysis, Tarailis et al. suggested that microstates A and B have been associated with sensory processing (microstate A: audition, arousal; microstate B: vision), microstates C and F with personally significant information processing, microstate D with attention-related processing, microstate E with interoceptive and emotional information processing, microstate G with somatosensory network (Tarailis et al., 2023). Several studies have characterized brain microstate changes from wakefulness to sleep (Bréchet et al., 2020; Brodbeck et al., 2012; Diezig et al., 2022; Wiemers et al., 2023). However, specific components of consciousness during REM sleep, such as emotions, visual perception, executive function, and metacognition, have not been clarified.

Therefore, this study aims to determine brain network activity associated with the conscious experience during lucid dreaming. Based on the functions of different microstates and the characteristics of lucid dreaming, we have a strong hypothesis that the presence of visual-related microstate B, self-related microstate C, cognitive control microstate D, and interoceptive-related microsite E will have a difference between lucid REM sleep and non-lucid REM sleep. We have a weak hypothesis about microstate A. We did not expect significant changes in microstates F and G. To test these hypotheses, we extracted the EEG data from non-lucid REM sleep and lucid REM sleep and contrasted them within the subject for the microstate analysis.

## Materials and Methods

### Participants

Nine proficient lucid dreamers (5 men and 4 women) participated in the study (age: 20–48 years). Participants were selected for high self-reported frequency of lucid dreams, with a demonstrated ability to successfully have lucid dreams in a sleep laboratory setting. All participants had no history of neurological disorder and had normal or corrected-to-normal vision. Signed informed consent was obtained from all participants before the experiment, and ethical approval was obtained from the Stanford University Institutional Review Board. The protocol was conducted in accordance with the declaration of Helsinki.

### Procedure

Participants arrived at 9:00 pm. After the EEG setup, participants were briefed on the lucid dreaming task by the experimenter. In addition to having lucid dreams, participants were instructed to perform the different lucid dream tasks (eye smooth tracking or hand clenching) in lucid dreams, as accurately as possible during the lucid dream. They were specifically trained to perform a specific left-right-left-right-center eye movement (LRLR), a standardized method for participants to communicate their lucid dreaming (La Berge et al., 1981). They were instructed to signal the beginning and ending of a lucid dream and the initiation and completion of each task trial during lucid dreaming (Fig. 1). Under the experimental constraints, we obtained as many trials from participants as possible.

**Figure 1.**
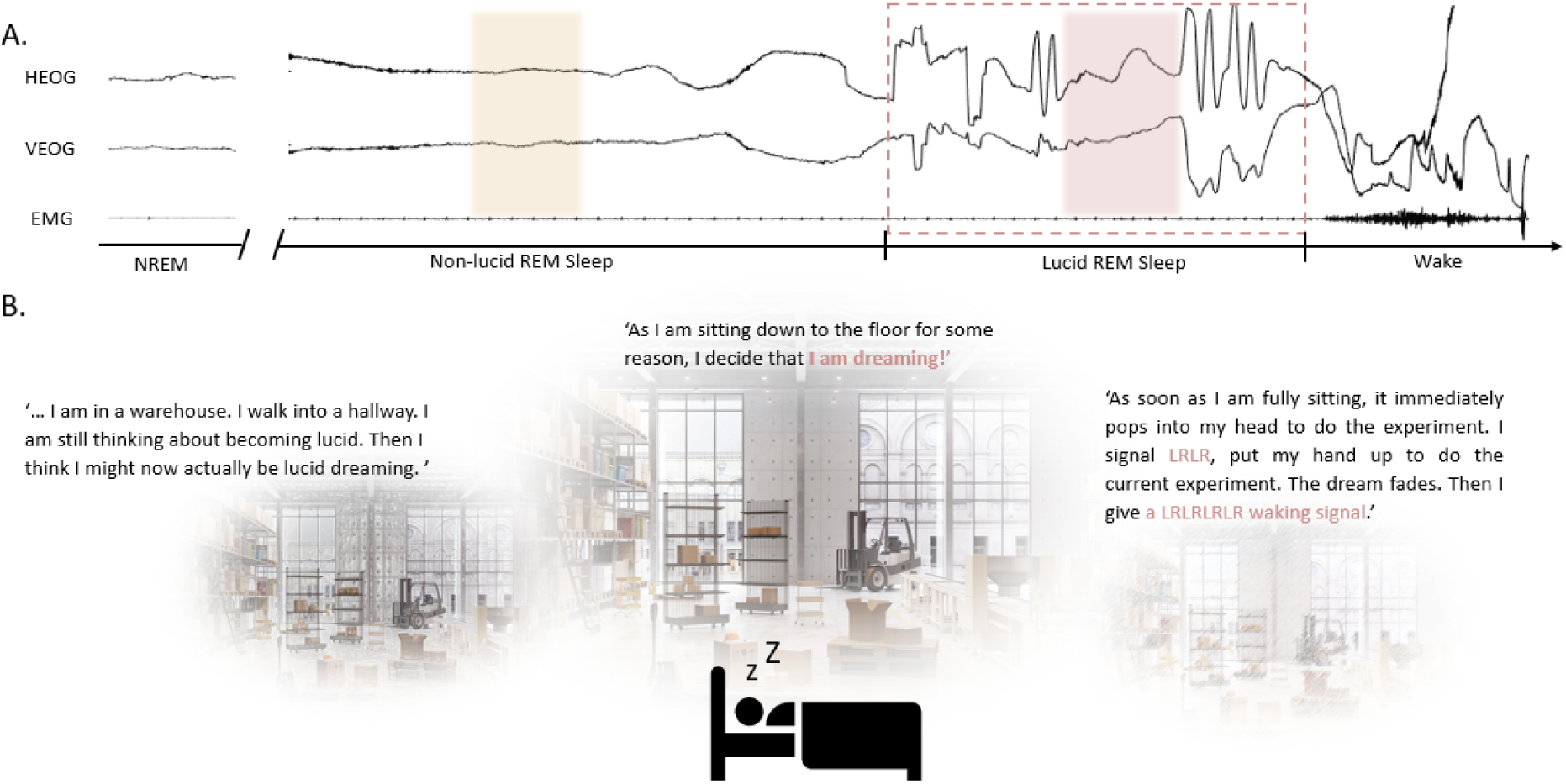
A lucid REM sleep dreaming from a single participant. A. Three channels of physiological data (HEOG, VEOG, and EMG) are shown during an initial period of NREM, REM period (90 seconds before lucid dreaming onset), transition to lucid REM sleep, and awakening. Data was bandpass filtered by 0.1 Hz. The dashed line shows a period of lucid dreaming: The participant reported becoming lucid in the dream and making an LRLR signal; when he was ready to wake up, he gave another LRLRLRLR signal. Red squares show the extracted artifact-free EEG data from lucid REM sleep. Yellow squares show the extracted artifact-free EEG data from non-lucid REM sleep. B. Dream report from this participant (see Supplementary Note 1).

After participants woke up from a lucid dream, they were instructed to write about their dream experiences in as much detail as possible. Subsequently, they filled out the Subjective Experiences Rating Scale (SERS) (Kahan, 1994), a questionnaire used to assess nine specific: sensory qualities, affective qualities, cognition, recall, sleepiness, general reality orientation, point of view, dreaming, and metacognition. Unfortunately, some of the questionnaires were missing, so we did not do further analysis with these questionnaires.

### EEG recording

Participants spent between 1 and 32 nights in the lab with continuous EEG recording. EEG signals were recorded by Neuroscan 32-channel system (Neuroscan, El Paso, TX, USA), with 29 EEG electrodes placed according to the international 10-20 system (Jasper, 1958), a submental electromyogram (EMG) electrode, and vertical and horizontal electrooculogram (EOG) electrodes. The sampling rate was 250 Hz. All electrodes were applied with Grass EC2 electrode paste and covered by a white net.

### EEG preprocessing

#### EEG Data Selection

The original data set recorded included 105 lucid dream recordings from 9 participants. These lucid dreams were validated through three different criteria (Erlacher & Stumbrys, 2020), and should meet all three criteria: (1) the participant’s subjective self-rating of lucidity; (2) the dream report rated by an external judge as having either possible or clear signs of lucidity; (3) the participant performs LRLR eye signaling, which can be unambiguously identified on the EEG recording by the research team.

From the 105 lucid dream recordings, each EEG recording was manually inspected to determine its eligibility for inclusion in the EEG microstate analysis. The inclusion criteria were as follows:1) clear visible eye-signaling (LRLR) at lucidity onset, 2) lucid dream recording segment with a duration longer than 6 seconds, and 3) minimal artifacts, such as muscle activity (myogenic artifacts), eye movements (ocular artifacts), and external interference. Lucid dream segments (lucid REM) containing onsets and offsets marked by LRLR signals were extracted from the EEG data. Any segments with significant ocular and muscular artifacts were manually excluded from each recording night. Then, an equivalent duration of segments (non-lucid REM) was extracted from the 90-second window of REM sleep preceding the lucidity onset from the same recording night (Fig. 1B).

After the selection, 39 recording nights from 8 participants met the inclusion criteria and were included for further analysis. Lucid-REM sleep (mean length 29.3±21.1 seconds) and non-lucid REM sleep (mean length 29.5±18.1 seconds) segments from each recording night were extracted for EEG microstate analysis (see Supplementary Table 1).

#### EEG Preprocessing

EEG data was analyzed using EEGLAB V2023.0 (Delorme & Makeig, 2004) and MATLAB R2022b (Mathworks Inc. Natick, MA, USA). Bad channels were identified by visual inspection and interpolated using spherical splines. EEG data were bandpass filtered between 0.5 Hz and 70 Hz, and notch filtered between 58Hz and 62Hz. Data were re-referenced to the average reference. Independent component analysis (ICA) was computed to remove ocular, muscular, and electrocardiographic artifacts. Finally, the remaining segments presenting any physiological or technical artifacts were removed by visual inspection.

### Microstate analysis

#### Identifying Individual Microstate Maps

The remaining data were filtered between 2 and 20 Hz. EEG microstate analysis was performed using MICROSTATELAB, an EEGLAB toolbox for resting-state microstate analysis (Nagabhushan Kalburgi et al., 2023). First, the Global Field Power (GFP) was computed for each sample over time. Topographies at the local maxima of GFP were clustered for each condition (lucid-REM sleep and non-lucid REM sleep) separately according to a fixed range of cluster numbers (4–7) using a modified k-means algorithm. The polarity of the topographies was ignored. Subsequently, the *condition mean* microstate maps were identified for the two conditions across individual microstate maps, and the *grand mean* microstate map was averaged by two *condition mean* microstate maps. Then, the *grand mean* microstate map was sorted based on published template maps (Custo et al., 2017), followed by sorting individual and *condition mean* maps by the *grand mean* map.

#### Microstate class solution

After cluster analysis, we got 4-7 class solutions. To determine the optimal number of microstate classes, we did our analyses following the logic as below: Based on the literature on the function of microstates and the characteristics of lucid dreaming, we had a strong hypothesis that the presence of visual-related microstates B, self-related microstate C, attention-related microstate D, and interoceptive-related microsites E will have a difference between lucid REM sleep and non-lucid REM sleep. We had a weak hypothesis about the emotional-related microstate A. We did not expect changes regarding microstates F and G. Thus, we chose the microstate class number by selecting the smallest set of grand mean microstate maps containing microstate classes A, B, C, D, and E.

#### Microstate Feature Extraction

To optimize the comparability of the microstate characteristics extracted (Nagabhushan Kalburgi et al., 2023), all initial individual electric potential field maps were assigned to the grand mean map, which guaranteed a consistent assignment across participants, resulting in a continuous sequence of microstate maps for each individual. Then, features of these identified microstates were extracted. These microstate dynamics features included the average duration in milliseconds (Duration), the average number of occurrences per second (Occurrence), the average coverage of the EEG signal in percent for each microstate class (Coverage), and finally, the explained variance for each microstate class (ExpVar).

### Statistical analysis

In order to assess potential associations between distinct microstate networks and specific aspects of lucid REM and non-lucid REM sleep, we performed separate linear mixed models for dependent variables of duration, occurrence, coverage, ExpVar. The condition (lucid-REM sleep and non-lucid REM sleep) and microstate class were used as fixed factors. A by-subject random intercept was added to represent between-person variability and account for the unbalanced data structure (Bates et al., 2015). We analyzed the data using RStudio 2023.6.1^1^, fitting all models with the package lme4 (Bates et al., 2015). ANOVA tests were performed to test each linear mixed-effects model for significance, and post hoc pairwise comparisons were conducted using the ‘emmeans’ (version 1.8.7) function in the RStudio package with Tukey correction. Type III tests of fixed effects were assessed using Satterthwaite’s method. In addition, to explore whether there are systematic differences in the spatial distribution of one or more microstate classes between different conditions, we compared the topography of the mean maps of the two conditions using topographic analysis of variance (TANOVA) (Nagabhushan Kalburgi et al., 2023).

### Meta-microstate Analysis

In previous studies, assessing microstate map similarity across studies typically relied on visually comparing topographies with previously published figures, which is imprecise and lacks a quantitative description. Also, the inability to search literature based on spatial topographies instead of microstate labels increases the risk of underestimating associations between studies (Koenig et al., 2023). To objectively relate our findings to the existing empirical evidence on EEG resting state microstates, we performed a meta-microstate analysis on our six class microstates using the MATLAB applications: MSTemplateEditor and MSTemplateExplorer (Koenig et al., 2023) (https://github.com/ThomasKoenigBern/MS-Template-Explorer). These apps relate microstate findings across studies by comparing the topography similarity, irrespective of the number of classes and labels chosen by the different research groups. We first added the microstate template maps from our data (grand mean microstate maps) using the MSTemplateEditor app. Subsequently, we assessed the topographical similarities between our microstate template maps and those from other studies in the database, generating a similarity matrix with the MSTemplateExplorer app. In our analysis, we selected empirical findings associated with microstate maps from the database that exhibited a topography similarity of over 65% with our template maps and were in some conceptual connection to our hypotheses. A discussion of the association of these empirical findings with our findings was conducted in descending order of topographic similarity.

## Results

### Microstate class solution

We first identified the microstate class solutions based on our hypothesis. In this study, The 4 class *grand mean* solution identified microstate classes C, D, E, F, and the 5 class *grand mean* solution identified microstate classes A, B, C, D, F, did not show all of the maps to test our strong hypotheses. The 6-class solution identified microstate classes A, B, C, D, E, and G, thus containing all classes we were interested in. It was, therefore, considered adequate for our purposes and maintained for further analysis (Figure 2). The 7 class solution (representing microstates A, B, C, D, E, F, G) only added a class (microstate F) that we had no hypothesis about and was therefore not considered further.

**Figure 2.**
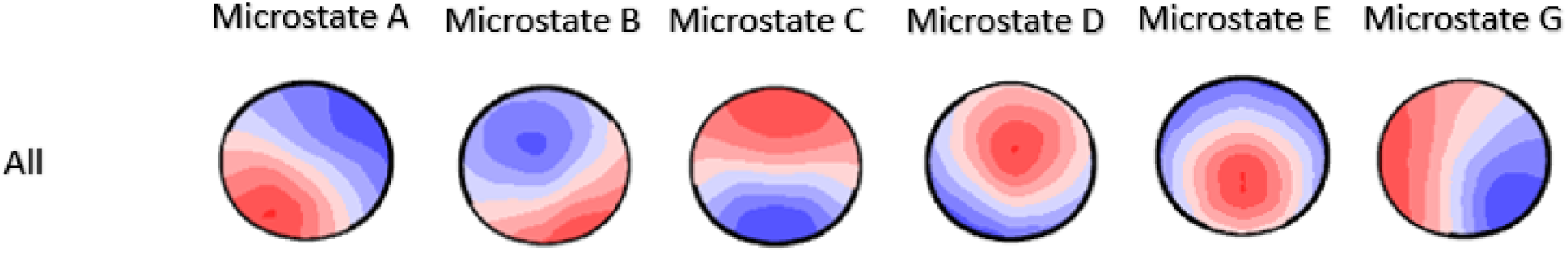
The six microstate topographies averaged across participants and conditions.

### TANOVA

To confirm the 6-class solution can demonstrate the global network changes in both conditions, we did a TANOVA on the microstate maps. There were no significant differences between microstate class maps of different conditions in the 6-class solution (all p-values above 0.1). This result indicates that generators of the maps of non-lucid REM sleep and lucid REM sleep were similar. Therefore, the parameters of these six microstate maps were considered sufficient to describe global network changes during both non-lucid REM sleep and lucid REM sleep. The resulting *grand mean* microstate maps of the 6-class solution are shown in Fig. 2.

#### Modeling of Microstate Parameters

Next, we observed differences between conditions in the temporal dynamics of the sixmicrostates (see Figure 3). The mixed models computed for the different microstate parameters yielded significant effects, including the factor condition for microstate coverage, duration, occurrence, and explained variance. The findings are visualized in Fig. 3 and briefly reviewed below.

**Figure 3.**
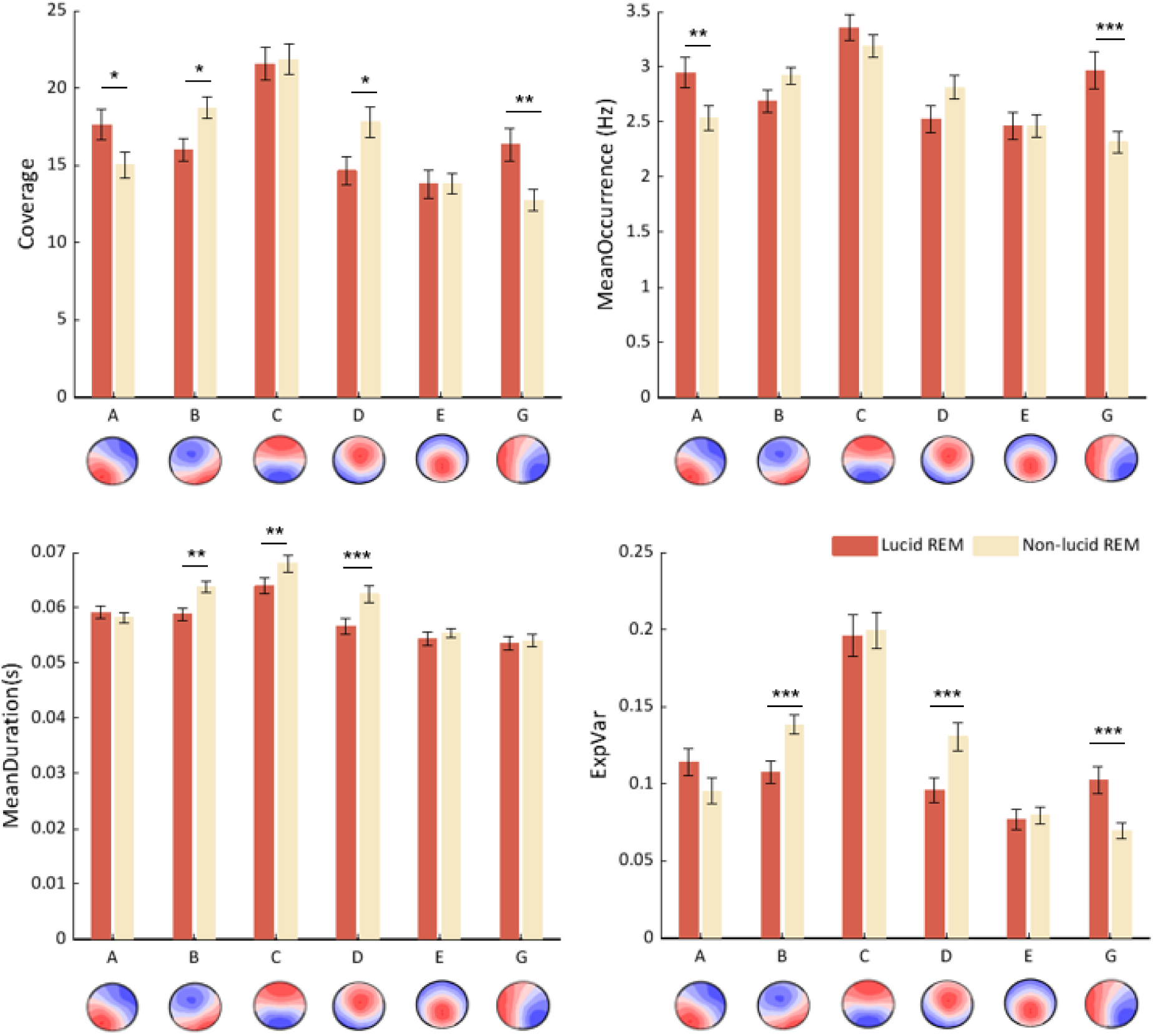
The mean and standard error of coverage (A), occurrence (B), duration (C), and Explained Variance (D) for each microstate class in both conditions. Error bar was calculated by Standard Error.

### Coverage

The results showed a significant interaction effect between microstate classes and condition (non-lucid REM vs. lucid REM) (F_(5, 456)_ = 5.07, *p* < 0.001). The conditional t-tests showed that coverage of microstate A (t(453) = 2.179, *p* <0.05) and E (t(453) = 3.031, *p* <0.01) were significantly higher during lucid REM sleep than non-lucid REM sleep, whereas microstate B (t(453) = 2.179, *p* <0.05) and D (t(453) = 2.534, *p* <0.05) were significantly higher during non-lucid REM sleep than lucid REM sleep. These results indicate that, compared to non-lucid REM sleep, the brain spent more time in neural networks represented by microstates A and E during lucid REM sleep and less time in neural networks represented by microstates B and D.

### Occurrence

The results showed a significant interaction effect between microstate class and condition (F_(5, 456)_ = 5.15, *p* < 0.001). There was no significant effect for the condition (*p*=0.056). In the subsequent conditional t-tests, the results showed that the microstate A (t(453) = 2.642, *p* <0.01) and E (t(453) = 4.133, *p* <0.0001) had a significantly higher frequency of activating during lucid-REM sleep than during non-lucid REM sleep.

### Duration

The results indicated a significant main effect for condition (F_(1, 443.45)_ = 15.61, *p* < 0.001) and a significant interaction effect between microstate class and condition (F_(5, 443.45)_ = 2.63, *p* < 0.05). In the follow-up t-tests, microstates of class B (t(450) = 2.988, *p* <0.01), C (t(450) = 2.598, *p* <0.01), and D (t(450) = 3.473, *p* <0.001) were found to last significantly longer during lucid-REM sleep than during non-lucid REM sleep, indicating that these microstates were more stable during the non-lucid REM sleep.

### Explained Variance

There was a significant interaction effect between microstate class and condition (F_(5, 452.72)_ = 5.17, *p* < 0.001). No significant main effect for the condition was found (*p*=0.86). The t-test results showed that the *Explained Variance* of microstate B (t(452) = 2.595, *p* <0.01) and D (t(452) = 2.926, *p* <0.01) were significantly higher during lucid-REM sleep than during non-lucid REM sleep, whereas the *Explained Variance* of microstate G (t(452) = 2.865, *p* <0.01) was lower during lucid-REM sleep than non-lucid REM sleep.

#### Cognitive function of microstates

To associate cognitive functions between different microstates more objectively and systematically, we performed a topographic similarity analysis between our six class microstates (See Methods) and a database of published microstate maps. Empirical findings associated with microstate maps were primarily chosen from studies on consciousness-related topics that included a within-subject variation of state (Table 1).

**Table 1.** Topographical similarities between our microstate template maps and those from the consciousness-related studies in the database.

| Study | Topic | MS A | MS B | MS C | MS D | MS G |
| --- | --- | --- | --- | --- | --- | --- |
| Lucid-REM v.s. nonlucid-REM |  | Lucidity during REM ↑ | Lucidity during REM ↓ | Lucidity during REM ↓ | Lucidity during REM ↓ | Lucidity during REM ↑ |
| (Custo et al., 2017) | Source Localization of Microstates | 94.6%<br>BA41 (PAC), BA22 (Wernicke), BA19, Left insular cortex ↑ | 77.5%<br>Cuneus, BA17 BA18, BA8 ↑ | 91.7%<br>PCC&Precuneus, left angular gyrus ↑ | 71.7%<br>BA40, BA13, Right middle and superior frontal gyri ↑ | 83.7%<br>Left middle frontal gyrus, Dorsal part of the anterior cingulate<br>Cuneus, extending to the PCC, Thalamus ↑ |
| (Britz et al., 2010) | BOLD correlates | 84.5%<br>Bilateral superior and middle temporal gyri, Left middle frontal gyrus ↓ | 67.0%<br>Bilateral inferior occipital gyri<br>Bilateral cuneus<br>Left lingual gyrus<br>Middle occipital gyrus ↓ | 91.7%<br>ACC, Cingulate gyrus medially ↓ | 78.7%<br>Right superior and inferior parietal lobules superior and middle frontal gyrus ↓ | 76.5%<br>Bilateral inferior occipital gyri<br>Bilateral cuneus<br>Left lingual gyrus<br>Middle occipital gyrus ↓ |
| (Croce et al., 2018) | Magnetic stimulation | 97.4%<br>Stimulated rAG↓<br>95.6%<br>Stimulated rIPS ↑ | 81.1%<br>Stimulated rAG ↓ | 97.6%<br>Stimulated TPJ -- | 87%<br>Stimulated rIPS ↓ | 91.1%<br>Stimulated AG topograph changed |
| (Bréchet et al., 2020) | Dreaming |  | 82.0%<br>Slowwave sleep vs wakeful resting↑<br>Dream experience↑ | 63.5%<br>Slowwave sleep vs wakeful resting↑<br>Dream experience↑ | 81.5%<br>Slowwave sleep vs wakeful resting↑<br>Dream experience↑ |  |
| (Diezig et al., 2022b) | Dream-like experience |  |  |  |  | 88.4%<br>Hypnagogic state ↓ |
| (Denzler et al., 2024) | Dream-like experience in VR | 68.7%<br>Bizarr vs normal VR environment (trend) ↓ |  | 77.4%<br>Bizarr vs normal VR environment ↑ | 75.5%<br>Bizarr vs normal VR environment (trend) ↓ |  |
| (ZanESCO, Denkova, et al., 2021) | Mind wandering |  |  | 93.8%<br>Mind wandering ↑ |  |  |
| (Tomescu et al., 2022) | Spontaneous thought | 84.6%<br>Planning, verbal thoughts↑<br>High control task ↑ | 85.9%<br>High control task ↑ | 90.1%<br>Self and Somatic Awareness↓<br>Planning, Comfort ↑<br>High control task ↑ | 63.4%<br>High control task ↑ | 91.7%<br>Planning, verbal thoughts↑ |
| (Tarailis et al., 2021) | Self-Reported Spontaneous Thoughts |  | 83.5%<br>Thinking about self↑ | 91.6%<br>Feeling of comfort↓ | 93.9%<br>Self↓ | 86.2%<br>Comfort↑ |
| (Schiller et al., 2021) | Acute alcohol intoxication |  | 78.0%<br>Intoxicated↑ | 95.4%<br>Intoxicated↓:<br>Transition C →B,<br>C→D↓<br>Somatic Awareness↓ |  | 71.5%<br>Intoxicated↑ |

\* '--' There were no statistically significant findings on this microstate map; '↑' Activity of microstate increased; '↓' Activity of microstate decreased. The findings in the blue regions exhibited a similar activity trend to what we observed in our study, while yellow exhibited the reverse trend. The white region exhibited the topographic similarity between our microstate map and the map from the database was below 65%, so we did not get related findings for further analysis.

Based on the empirical findings associated with similar microstate maps, we mainly focused on the microstates in which we got significant results.

*Microstate A.* According to the findings associated with the maps selected based on spatial similarity, microstate A might connect with arousal. Studies using source imaging techniques or combined EEG-fMRI reported activity for microstate A in areas partially overlapping with the left middle and superior temporal lobe, primary auditory cortex, and left insular cortex (Custo et al., 2017) (94.6%), as well as the bilateral superior, middle temporal gyri and left middle frontal gyrus (Britz et al., 2010) (84.5%).

*Microstate B.* microstate B was primarily related to visual processing. For example, the source localization study on microstate B with 77.5%) showed very strong activity in visual areas: in the left and right occipital cortices (cuneus), including Brodmann areas 17 and 18 (primary visual cortex) (Custo et al., 2017).

*Microstate C.* Based on previous literature, the function of microstate C seems somewhat ambiguous, showing characteristics of both a task-negative network and a control-related network. For instance, the presence of microstate C was associated with mind-wandering (93.8%) (Zanesco et al., 2021), somatic awareness (91.6%) (Tarailis et al., 2021), and showed overlap with DMN regions (91.7%) (Custo et al., 2017). However, in our study, stronger evidence with higher spatial map similarity supports its association with control-related network functions, suggesting that microstate might be more involved in cognitive processes related to monitoring. For example, Schiller et al. reported that alcohol reduced the presence of microstate C (95.4%), which was associated with a decline in higher-order processing of internal information and cognitive control. (Schiller et al., 2021). Additionally, source localization evidence also showed that microstate C overlapped with the salience network (91.7%) (Custo et al., 2017).

*Microstate D.* From source localization and combined EEG-fMRI studies, microstate D (78.7%, 71.7%) generally overlapped with the frontoparietal network (Britz et al., 2010; Custo et al., 2017). This was further supported by a brain stimulation study, which found that applying transcranial magnetic stimulation (TMS) over the right intraparietal sulcus (IPS), a node of the dorsal attention network, could alter the presence of microstate D (87%) (Croce et al., 2018). The presence of similar microstate maps is also associated with high cognitive control (63.4%) and externally orientated processing (93.9%) (Tarailis et al., 2021; Tomescu et al., 2022).

*Microstate G.* In our study, microstate G proved to be associated with Default Mode Network (DMN). For example, studies on spontaneous thought during wakefulness showed that the appearance of similar maps was positively correlated with self-related thoughts(91.7%) (Tomescu et al., 2022) and self-comfort (86.2%) (Tarailis et al., 2021). TMS study (91.1%) (Croce et al., 2018) and source localization study (83.7%) (Custo et al., 2017) also identified similar maps associated with the DMN.

## Discussion

The results of our study show that microstates B, C, and D were less present during lucid REM sleep, while microstates A and G dominated during non-lucid REM sleep. Considering the inhibitory function of microstates reported in studies of dreaming and dream-like experiences (Bréchet et al., 2020; Deolindo et al., 2021; Diezig et al., 2022), along with the characteristics of lucid dreaming mentioned earlier, lucid REM sleep was associated with increased visual processing, metacognition, and volitional control; decreased emotional arousal and DMN activity. The pattern of our results confirmed an inverse relationship between microstates and functional correlates, which suggested that the activity of microstates showed the deactivation of related brain networks during REM sleep.

### Visual processing during REM sleep

Lucid dream is generally more perceptually vivid than the average nonlucid dream (LaBerge & DeGracia, 2000). Besides self-reports, this phenomenon was supported by neuroimaging measuring showing greater activation of visual areas during lucid REM sleep compared to non-lucid REM sleep (Dresler et al., 2012). A recent study also showed that the vividness of the experience can be affected by volition control during lucid dreaming (Raduga et al., 2020). In sum, converging evidence suggests lucid dreaming might have more intense visual activation than non-lucid REM sleep. Our study found that the presence of microstate B, decreased during lucid REM sleep compared to non-lucid REM sleep. Based on the assumption of more active visual processing during lucid dreaming, and in line with the potential inhibitory function of microstates, our results suggest that the reduction of microstate class B is indeed a correlate of intense visual processing during lucid dreaming. This fits the findings from a hypnagogic study: the visual character of dream-like experiences was related to the decreased presence of a microstate associated with higher-order visual areas (Diezig et al., 2022). In sum, our study supports the hypothesis that the decreased presence of microstate B represents a more intense visual perception during lucid REM sleep compared to non-lucid REM.

Notably, an intriguing finding is that our microstate B topography had a high similarity with a microstate (83.5%) related to self-reflection in Tarailis et al., (2021). Similar experiences are typical during lucid dreaming, with the activation of the *observing-monitoring-reflecting self* (Kahan & LaBerge, 1994). Again, assuming an inhibitory effect of microstates, the decreased presence of microstate B might also suggest that self-reflection is more dominant during lucid REM sleep. However, given the limited evidence regarding the relationship between microstate B and self-reflection in previous studies, we cannot conclusively confirm the self-reflective function of our microstate map. Future studies can clarify this relationship further.

### Volitional dream control during REM sleep

During lucid dreaming, dreamers can control their own behaviors within the dream but also shape the dream itself volitionally (LaBerge & DeGracia, 2000; Stumbrys & Erlacher, 2017). Such dream control ranges from simple tasks like performing reality checks or flying to more complex actions, such as executing prearranged tasks like clenching their hands, performing mathematical calculations, or even communicating with researchers during the lucid dream (Erlacher et al., 2014; Konkoly et al., 2021; Peters et al., 2024). In contrast, non-lucid dreams almost lack this kind of volitional dream control (Voss et al., 2013). Thid can be confirmed by the decreased presence of cognitive control-related microstate D during lucid REM sleep compared to non-lucid REM sleep. Aligning with the inhibitory effect of microstate, this reduction reflects increased cognitive control and frontoparietal activities typical in lucid dreaming. This result is also consistent with previous studies, which found an increased BOLD signal (Dresler et al., 2012b) and a greater beta-1 activity (Holzinger et al., 2006) in the parietal regions during lucid dreaming compared to non-lucid dreaming. A recent NREM sleep study found that a similar microstate map (81.5%) was more present during sleep compared to wakefulness, indicating deactivation of frontal brain areas during sleep (Bréchet et al., 2020). These evidence supports that the decreased presence of microstate D represents a heightened dream control during lucid REM sleep compared to non-lucid REM. Considering the limited number of channels in our study, we were unable to precisely estimate the sources contributing to our microstate D. Further research with high-density EEG and source analysis could help clarify the specific neural sources involved in cognitive control during lucid dreaming.

### Activation of default mode network during REM sleep

DMN is the likely neural correlate of dreaming: where externally generated experiences and voluntary control were reduced, internally generated and self-focused experiences became more central, leading to increased DMN activity (Domhoff & Fox, 2015; Fox et al., 2013). It generally showed anticorrelations with the ‘task-positive network’(Raichle, 2015). Following this logic, compared to non-lucid dreaming, dreamers partly regained sensory awareness and voluntary control during lucid dreaming, which increases the involvement of networks related to external attention and control and reduces the recruitment of the DMN. Our results are consistent with this notion, showing that the presence of microstate G was dominant during lucid REM sleep compared to non-lucid REM sleep. Aligning with the inhibitory effect of microstates, this finding indicated that the DMN was more activated during non-lucid REM sleep compared to lucid REM sleep. Similarly, Diezig et al. found that the decreased presence of similar map (88.4%) correlated with increasing hypnagogic experiences (Diezig et al., 2022). The brain areas underlying this map were shown in right-lateralized activity in the middle and inferior temporal gyrus, which partly overlapped with the subsystem of the DMN reported in Andrews-Hanna et al., 2010 study. Therefore, we suggested that the increased presence of microstate G represents a deactivation of DMN during lucid REM sleep compared to non-lucid REM.

### Activated metacognition during REM sleep

The salience network is associated with performance monitoring (Oliveira et al., 2007) and plays an important role in metacognition (Ham et al., 2013). Recently, Asakage et al. found that the salience network was activated during self-recognition from both first-person and third-person perspectives (Asakage & Nakano, 2023). These functional properties align well with the experience of lucid dreaming: dreamers have metacognitive insight into the fact that they are dreaming and can exert volitional control over their experiences. In contrast, non-lucid dream generally exhibits decreased metacognitive processing, often characterized by deficits in self-monitoring and confusion between internally and externally generated experiences (Perogamvros et al., 2017). Based on these findings, we expected that metacognition would be more dominant during lucid REM sleep.

Our result showed a decreased duration of microstate C during lucid REM sleep compared to non-lucid REM sleep. Again, considering the inhibitory effect of the microstate, the salience network might be more recruited during lucid REM sleep. In lucid dreaming, participants partly regain sensory awareness and voluntary control, leading to increased conflicts between self-generated and externally generated experiences. As a result, the salience network might be more recruited to monitor and resolve these conflicts, leading to its prolonged activation. Similarly, a recent study found that increased contribution of microstate C’ (77.45% similarity) was associated with suspension of disbelief in VR context, such as the intention to disregard mismatches between the current sensory experience in the virtual reality and prior expectations (Denzer et al., 2024). Therefore, in our study, the decreased presence of microstate C represents a heightened metacognition during lucid REM sleep compared to non-lucid REM.

### Emotion during lucid dreaming

For emotional arousal, the increased presence of microstate A during lucid REM sleep compared to non-lucid REM sleep might suggest that the emotional-related network is less active during lucid dreaming. The intense emotions often experienced in non-lucid dreams can be explained by the activation of the limbic system (Braun et al., 1997; Maquet et al., 1996). During lucid dreaming, the dreamer’s awareness of being in a dream and their ability to monitor the dream state may contribute to modulation or reduction in emotional intensity (de Macêdo et al., 2019; Tzioridou et al., 2022). Our findings on reduced emotional arousal provide a preliminary indication of differences in emotional processing between lucid and non-lucid REM sleep. Future studies should investigate these differences further.

### Summary

According to the definition of lucid dreams, dreamers can either actively influence the events of the dream or passively observe the progression of the dream (Stumbrys et al., 2013). In our study, we mixed these two conditions. Participants were instructed to perform eye movements to signal the lucidity onset and offset during lucid REM sleep; in some lucid dreams, participants were capable of using eye movements to initiate and complete a task during lucid REM sleep, while others were not. It is possible that the presence of the related microstate maps still partially reflects task-related execution rather than lucidity itself. Future studies need to explore the neural correlates of active and passive lucid dreams in more detail. Another limitation is the imbalance in the contribution of data from individual participants (Supplementary Table 1). Therefore, findings should be interpreted cautiously and carefully generalized across the entire sample.

In conclusion, Our findings demonstrate that EEG microstates can reveal distinct network dynamics in lucid and non-lucid REM sleep, linking specific global brain networks to variations in specific consciousness content. Future research could further investigate how these networks interact during different dream experiences.

## Acknowledgment

This work was supported by the Swiss National Science Foundation (SNSF), no. 10001CM_201074. We would like to express our gratitude to all the lucid dreamers for their time and effort, which made this study possible. We also thank Sarah Diezig for assisting with the data analysis and Keith Garcia for project organization and data collection.

## Footnotes

1 Posit team (2023). RStudio: Integrated Development Environment for R. Posit Software, PBC, Boston, MA. URL http://www.posit.co/.

## Notes

### Competing Interest Statement

The authors have declared no competing interest.

