## Supplementary Information for "EEG Microstates reveal distinct network dynamics in lucid and non-lucid REM sleep"

### Supplementary Table 1. Microstate analysis with all recording nights.

| DataSet | Subject | Condition | EEG length | Condition | EEG length | Task | Channel | Questionnaire | Dream report |
|---|---|---|---|---|---|---|---|---|---|
| Sub01_1 | Sub01 | Lucid REM | 56.52 | non-lucid REM | 48.69 | None | 29 | Yes | Yes |
| Sub02_1 | Sub02 | Lucid REM | 49.29 | non-lucid REM | 54.75 | None | 29 | Yes | Yes |
| Sub03_1 | Sub03 | Lucid REM | 24.75 | non-lucid REM | 29.2 | Clenching | 28 | No | Yes |
| Sub03_2 | | Lucid REM | 27.84 | non-lucid REM | 28.96 | Clenching | 29 | Yes | Yes |
| Sub03_3 | | Lucid REM | 14.35 | non-lucid REM | 26.34 | Clenching | 29 | Yes | Yes |
| Sub04_1 | Sub04 | Lucid REM | 47.12 | non-lucid REM | 40.37 | Circle | 29 | Yes | Yes |
| Sub04_2 | | Lucid REM | 18.06 | non-lucid REM | 20.12 | Circle | 29 | Yes | Yes |
| Sub05_1 | Sub05 | Lucid REM | 19.74 | non-lucid REM | 24.21 | Circle | 29 | Yes | Yes |
| Sub06_1 | Sub06 | Lucid REM | 41.44 | non-lucid REM | 42.96 | Circle | 29 | Yes | Yes |
| Sub06_2 | | Lucid REM | 41.65 | non-lucid REM | 42.71 | convergence | 28 | No | Yes |
| Sub07_1 | Sub07 | Lucid REM | 110.31 | non-lucid REM | 97.03 | None | 29 | Yes | Yes |
| Sub07_2 | | Lucid REM | 19.82 | non-lucid REM | 18.85 | Circle | 29 | Yes | Yes |
| Sub07_3 | | Lucid REM | 11.06 | non-lucid REM | 10.59 | None | 29 | Yes | Yes |
| Sub07_4 | | Lucid REM | 19.36 | non-lucid REM | 20.98 | Circle | 29 | Yes | Yes |
| Sub07_5 | | Lucid REM | 17.82 | non-lucid REM | 19.4 | Circle | 29 | Yes | Yes |
| Sub08_1 | Sub08 | Lucid REM | 6.31 | non-lucid REM | 11.1 | Circle | 29 | | Yes |
| Sub08_2 | | Lucid REM | 18.05 | non-lucid REM | 20.07 | Circle | 29 | | Yes |
| Sub08_3 | | Lucid REM | 22.55 | non-lucid REM | 21.69 | Circle | 29 | | Yes |
| Sub08_4 | | Lucid REM | 18.91 | non-lucid REM | 18.72 | Circle | 29 | | Yes |
| Sub08_5 | | Lucid REM | 32.27 | non-lucid REM | 29.29 | None | 29 | | Yes |
| Sub08_6 | | Lucid REM | 27.77 | non-lucid REM | 25.75 | Circle | 29 | | Yes |
| Sub08_7 | | Lucid REM | 11.04 | non-lucid REM | 11.07 | Circle | 29 | | Yes |
| Sub08_8 | | Lucid REM | 13.34 | non-lucid REM | 14.65 | None | 29 | | Yes |
| Sub08_9 | | Lucid REM | 51.4 | non-lucid REM | 45.61 | None | 29 | | Yes |
| Sub08_10 | | Lucid REM | 11.29 | non-lucid REM | 13.66 | None | 29 | | Yes |
| Sub08_11 | | Lucid REM | 11.67 | non-lucid REM | 18.88 | Circle | 29 | | Yes |
| Sub08_12 | | Lucid REM | 62.6 | non-lucid REM | 46.69 | Circle | 29 | | Yes |
| Sub08_13 | | Lucid REM | 61.01 | non-lucid REM | 56.85 | Circle+clenching | 29 | | Yes |
| Sub08_14 | | Lucid REM | 25.21 | non-lucid REM | 23.72 | Circle | 28 | | Yes |
| Sub08_15 | | Lucid REM | 19.91 | non-lucid REM | 17.59 | None | 28 | | Yes |
| Sub08_16 | | Lucid REM | 9.51 | non-lucid REM | 11.29 | None | 28 | | Yes |
| Sub08_17 | | Lucid REM | 7.74 | non-lucid REM | 10.35 | None | 29 | Yes | Yes |
| Sub08_18 | | Lucid REM | 61.63 | non-lucid REM | 63.81 | None | 29 | Yes | Yes |
| Sub08_19 | | Lucid REM | 7.52 | non-lucid REM | 10.79 | None | 28 | | Yes |
| Sub08_20 | | Lucid REM | 18.35 | non-lucid REM | 19.25 | convergence | 28 | | Yes |
| Sub08_21 | | Lucid REM | 31.83 | non-lucid REM | 33.12 | Line | 28 | | Yes |
| Sub08_22 | | Lucid REM | 21.92 | non-lucid REM | 25.63 | None | 28 | | Yes |
| Sub08_23 | | Lucid REM | 29.67 | non-lucid REM | 31.41 | None | 28 | | Yes |
| Sub08_24 | | Lucid REM | 43.68 | non-lucid REM | 43.18 | None | 28 | | Yes |

**Supplementary Note 1. Examples of participant reports during lucid REM sleep (showed in figure 1)**

From Sub08_17

I was having a dream that my family moved into a new house. My room was in the back, and my sisters' bedroom nearby. My brother's was toward the front. My parents and siblings were about ready to leave for church. I wasn't going because I didn't believe in attending church anymore.

I had a pile of dirty clothes lying in the middle of the floor. There were a lot of people there. I wondered what they would think about the dirty clothes. I began picking up the clothes and moving them to the laundry area. My parents walked out to leave. I wondered if my dad believed in God anymore or if he was just going to church for the hell of it.

I went to the front of the house where my brother's room was. I was looking for a better place to sleep. I saw my dad's car out front, a large deep-red metallic American car that he doesn't own in the waking world. I thought it wouldn't be a good place to sleep because it would be too noisy when the car was started in the morning.

I noticed the room was divided into two parts each as large or larger than mine. My mom came in and I asked her about the room. She became upset because she thought I was trying to get the room from my brother. Several times, I tell her, No, that I am not trying to. But, she doesn't believe me.

Then I am in this other scene where I am wandering around trying to become lucid. It seems there may be a part in the middle that I do not recall. I think that I should go lie down and relax and become lucid. Eventually, I lay down, decide to relax, and to forget all about lucid dreaming. I may have already been lying down in the dream equivalent of the lab booth or somewhere. In any case, I try to relax and fall asleep. I think all this occurred before I was in a warehouse.

Then, I am in a warehouse. I walk into a hallway. I am still thinking about becoming lucid. Then I think I might now actually be lucid dreaming. As I am sitting down to the floor for some reason, I decide that I am dreaming. As soon as I am fully sitting, it immediately pops into my head to do the experiment. I signal LRLR, put my hand up to do the current experiment. The dream fades. Then I give a LRLRLRLR waking signal.
